# Adipocyte Aldosterone in Obstructive Sleep Apnea

**DOI:** 10.1101/2025.01.07.631499

**Authors:** Yunzhou Dong, Josh Bock, Shahid Karim, Prachi Singh, Virend K. Somers

## Abstract

Aldosterone, a hormone synthesized by the adrenal glands, plays a pivotal role in regulating both blood pressure and electrolyte balance and may contribute to long term cardiovascular risk. Recent findings suggest that adipose tissue might serve as a source for aldosterone production and secretion. Patients suffering from obstructive sleep apnea (OSA) often exhibit elevated blood pressure levels and increased cardiovascular risk. Therefore, we sought to investigate whether increased aldosterone synthesis in fat tissue could contribute to the increase in hypertension and cardiovascular disease in OSA patients. To address this, we conducted a comparative analysis of adipocyte aldosterone level in OSA patients versus Age and BMI matched healthy individuals. Our data revealed no significant differences in aldosterone content between the two groups. Further examination involved measuring the expression of the key enzyme CYP11B2 responsible for aldosterone synthesis. We observed comparable expression levels of CYP11B2 in both the OSA and control groups. To corroborate these findings, we isolated stromal vascular fraction (SVF) from human fat tissues. The SVF cells were cultured in preadipocyte medium and subjected to normoxia and intermittent hypoxia (IH), mimicking OSA conditions as per established protocol. Subsequent analysis of aldosterone levels in cell lysates and conditioned medium revealed no significant differences. Similarly, *in vitro* examination of CYBP11B2 expression was not different in normoxic versus IH conditions. In conclusion, our study did not discern significant differences in adipocyte aldosterone production and secretion between patients with OSA and matched control subjects, nor in human preadipocytes under normoxic *versus* IH conditions.

## Introduction

Hypertension affects millions worldwide and is a significant risk factor for cardiovascular diseases (Wolk *et al*., 2003; Mills *et al*., 2020). Aldosterone, a hormone produced by the adrenal glands, plays a crucial role in regulating blood pressure and electrolyte balance (Pimenta & Calhoun, 2007; Gaddam *et al*., 2009; Ferreira *et al*., 2021; Leopold & Ingelfinger, 2023) and has been implicated in cardiovascular risk(Melby, 2002; Funder & Reincke, 2010; He & Anderson, 2013; Buffolo *et al*., 2022). Aldosterone primarily acts on the kidneys, where it promotes sodium reabsorption and potassium excretion (Muto, 1995; Epstein, 2001). Through its interaction with mineralocorticoid receptors (MR) in the distal tubules of the kidney, aldosterone increases the expression of sodium channels and sodium-potassium ATPases, leading to enhanced sodium retention and potassium excretion. Consequently, this results in increased extracellular fluid volume and arterial blood pressure (Fuller & Young, 2005).

In hypertensive individuals, dysregulation of the renin-angiotensin-aldosterone system (RAAS) may lead to excessive aldosterone production(Jia *et al*., 2018). Chronic activation of aldosterone receptors contributes to sodium retention, volume expansion, and vasoconstriction, thereby raising blood pressure (Ray *et al*., 2015). Moreover, aldosterone-induced endothelial dysfunction and vascular remodeling further exacerbate hypertension and promote end-organ damage (Ferreira *et al*., 2021).

Adipose tissue has emerged as a source for production of aldosterone (Briones *et al*., 2012; Dinh Cat *et al*., 2016; Kawarazaki & Fujita, 2016). Adipocytes express enzymes involved in aldosterone production, including aldosterone synthase (CYP11B2)(Dinh Cat *et al*., 2016), suggesting a local source of this hormone independent of adrenal glands. Furthermore, Briones et al reported that adipocytes produce aldosterone through calcineurin-dependent signaling pathways using 3T3-L1 cell line, human fat tissue and the db/db mouse model (Briones *et al*., 2012).

Understanding the role of adipose tissue-derived aldosterone has clinical implications (Sowers *et al*., 2009; Kawarazaki & Fujita, 2016). Individuals with obesity, metabolic syndrome, or insulin resistance exhibit dysregulated aldosterone production, predisposing them to hypertension (Sowers *et al*., 2009; Bothou *et al*., 2020).Targeting adipose tissue aldosterone synthesis or its downstream effects may offer novel therapeutic strategies for hypertension management (Taylor & Abdel-Rahman, 2009; Arendse *et al*., 2019; Ghatage *et al*., 2021; Desai *et al*., 2023; Cohen & Bress, 2024).

Adipose tissue-derived aldosterone may represent a novel paradigm in blood pressure regulation. Numerous studies have indicated that patients with OSA exhibit elevated blood pressure and body mass index (BMI) (Narkiewicz *et al*., 2005; Belyavskiy *et al*., 2019; Yeghiazarians *et al*., 2021). Additionally, obesity has been correlated with hypertension (Hsueh & Buchanan, 1994; Landsberg *et al*., 2013; Seravalle & Grassi, 2017; Litwin & Kulaga, 2021), suggesting a potential mechanism whereby adipose tissue in OSA patients could produce and secrete more aldosterone. To test this hypothesis, we analyzed aldosterone production and secretion from adipose tissue in OSA patients and matched control subjects, as well as *in vitro* studies of aldosterone secretion by preadipocytes exposed to normal oxygen levels compared to intermittent hypoxia (IH).

## Materials and methods

### Human subcutaneous abdominal fat tissues and cell lysis

Subjects: We studied 12 patients with OSA (2 female, 10 male) and 12 healthy control subjects (3 female, 9 male) matched for age and BMI. All control subjects underwent complete overnight polysomnography to confirm absence of OSA. This study was approved by the Mayo Clinic Institutional Research Board. All participants provided written informed consent prior to any study procedures.

Fat tissue preparation: Approximately 100mg of adipose tissue was weighed and finely ground in 1.5ml Eppendorf tubes containing a lysis buffer consisting of 50 mM Tris-HCl (pH 6.8), 6.8 M urea, 2% SDS, and 1X protease inhibitors cocktail (Roche), followed by 2x30 second sonication. Cell lysates were then centrifuged at 15,000xg for 15 minutes at 4°C, and the lipid layer at the top was meticulously removed. This process was repeated three times, or until the cell lysates appeared clear. Protein concentrations were determined using the BCA protocol as per the manufacturer’s instructions. Subsequently, cell lysates were utilized for Western blot analysis or aldosterone content assessment.

### Measurement of aldosterone content

To quantify aldosterone levels in fat tissue lysates, preadipocyte lysates, and conditioned medium, we adhered to the guidelines outlined in the aldosterone kit instructions provided by Eagle Company (cat# KLD31-K01). The following is a concise breakdown of the procedure: 1) Preparation of standard solutions containing known aldosterone concentrations for calibration, followed by the dilution of cell lysates to ensure that the aldosterone levels fall within the standard range. 2) Utilization of an ELISA kit to measure the optical density (OD) value at a wavelength of 450 nm. 3) Data analysis involved comparing the absorbance or signals obtained to a standard curve, while adjusting for dilution factors and subtracting the solvent reading to eliminate background interference.

### Isolation of human stromal vascular fraction (SVF) and preadipocyte culture

We implemented an SVF isolation procedure based on a previously established protocol, with minor modification (Hearnden *et al*., 2021). In brief, human abdominal fat tissues obtained from biopsies were finely minced using scissors and then placed in 15-ml tubes containing 5-6 ml of a 10 mg/ml collagenase type I (cat# LS004196, Worthington) enzymatic solution. The mixture was incubated for 45 minutes at 37°C in a slow rocker incubator. Following this, the solution was filtered through a 100 μm cell strainer (cat# 352360, Fisher Scientific) and then centrifuged at 800x g for 10 minutes at 4°C. The supernatant was aspirated, and the pellet was washed once with DMEM medium. Subsequently, the pellet was resuspended in preadipocyte culture medium (ZenBio Inc, cat# PM-1) supplemented with 10% FBS and antibiotics, and transferred to a 10 cm petri dish for culture. The medium was changed to fresh cell culture medium after 2-3 day of initial culture, with subsequent medium changes every 3 days.

### Intermittent hypoxia (IH)

For the *in vitro* cellular intermittent hypoxia (IH) model, we established preadipocyte culture models initially cultured under standard oxygen conditions (21% O_2_ and 5% CO_2_) and then in an intermittent hypoxia chamber, as described earlier (Polonis *et al*., 2020). Within the IH chamber, the cells underwent cycles comprising 30-minute intervals at 0.1% O_2_ and 5% CO_2_, followed by exposure to 21% O_2_ and 5% CO_2_ for 9 hours daily, over a period of 7 days. This experimental protocol was independently repeated more than three times to ensure reliability and reproducibility.

### Measurement of CYP11B2 expression by western blot

The western blot procedure, as previously reported (Dong *et al*., 2015; Dong *et al*., 2020), was employed for both fat tissue lysates and cell lysates. In brief, fat tissue samples were lysed using a buffer solution consisting of 50 mM Tris-HCl (pH 6.8), 6.8 M urea, 5% mercaptoethanol, 2% SDS, 1 mM PMSF, 1X protease inhibitors cocktail (Roche), 20 mM NaF, and 1 mM Na3VO4. The lysate underwent sonication on ice for 1 minute to enhance cellular protein extraction, repeating the process twice. Subsequently, the proteins were separated using 8.5% SDS– polyacrylamide gel electrophoresis and transferred onto a polyvinylidene fluoride membrane (Immobilon-P, Millipore-Sigma). Following transfer, the membranes were probed with specific antibodies, such as CYP11B2, and the protein bands of interest were visualized using SuperSignal™ West Pico PLUS Chemiluminescent Substrate (Thermo Fisher Scientific). Finally, the blot images were captured utilizing the Odyssey Fc Imager system (LI-COR).

### Determination of messenger RNA (mRNA) expression of CYP11B2 by qPCR

Quantitative real-time PCR analysis was conducted following a standardized protocol (Dong *et al*., 2023). Initially, total RNA was extracted using Trizol (Invitrogen), and its concentration was determined using a nanodrop spectrophotometer. Subsequently, cDNA was synthesized using the iScript™ cDNA Synthesis Kit (Bio-Rad). Specific oligonucleotide sequences were employed for the target human gene CYP11B2 (Forward: 5’- AGGCCTGAGCGGTATAATCC-3’; Reverse: 5’-AGTGTCTCCACCAGGAAGTG-3’), while β-actin served as the internal control (Forward: 5- CCCTGGAGAAGAGCTACGAG-3; Reverse: 5’-GGAAGGAAGGCTGGAAGAGT- 3’).

For the qPCR reaction, premixtures were obtained from Apenzybio (cat# CW2621F FastSYBR mixture). Real-time PCR was performed using the QuantStudio™ 5 Real-Time PCR System with the following cycling parameters: initial denaturation at 95°C for 10 min, followed by 40 cycles of denaturation at 95°C for 10 seconds and annealing/extension at 60°C for 1 min. The melting stage included steps at 95°C for 15 seconds, 60°C for 1 minute, and then holding at 95°C for 15 seconds. Fold change of mRNA can be determined by the formula r = 2^-ΔΔCt^ for relative quantification of gene expression.

### Statistical analysis

Statistical analysis was performed by Prism software v.10 (GraphPad). The data are presented as mean±SEM. Data analysis was carried out using Student *t* test. p<0.05 was considered a statistically significant difference.

## Results

### Demographic information - patients with obstructive sleep apnea and control subjects

Within the OSA group, there were 2 individuals with mild OSA (AHI 5-15), constituting 16.7%; 2 with moderate OSA (AHI 15-30), also 16.7%; and 8 with severe OSA (AHI >30), comprising 66.7%. All individuals in the control group had an AHI below 5.

As depicted in Fig. 1, the groups were well-matched in terms of age, sex, height, body weight (BW) and BMI (Fig. 1A, 1D to F). The primary differences lay in AHI (Fig.1B & C). Average AHI in the OSA group was 39.5, compared to 2.2 events/hour in the control group, p<0.0001 (Fig. 1B).

**Fig. 1.**
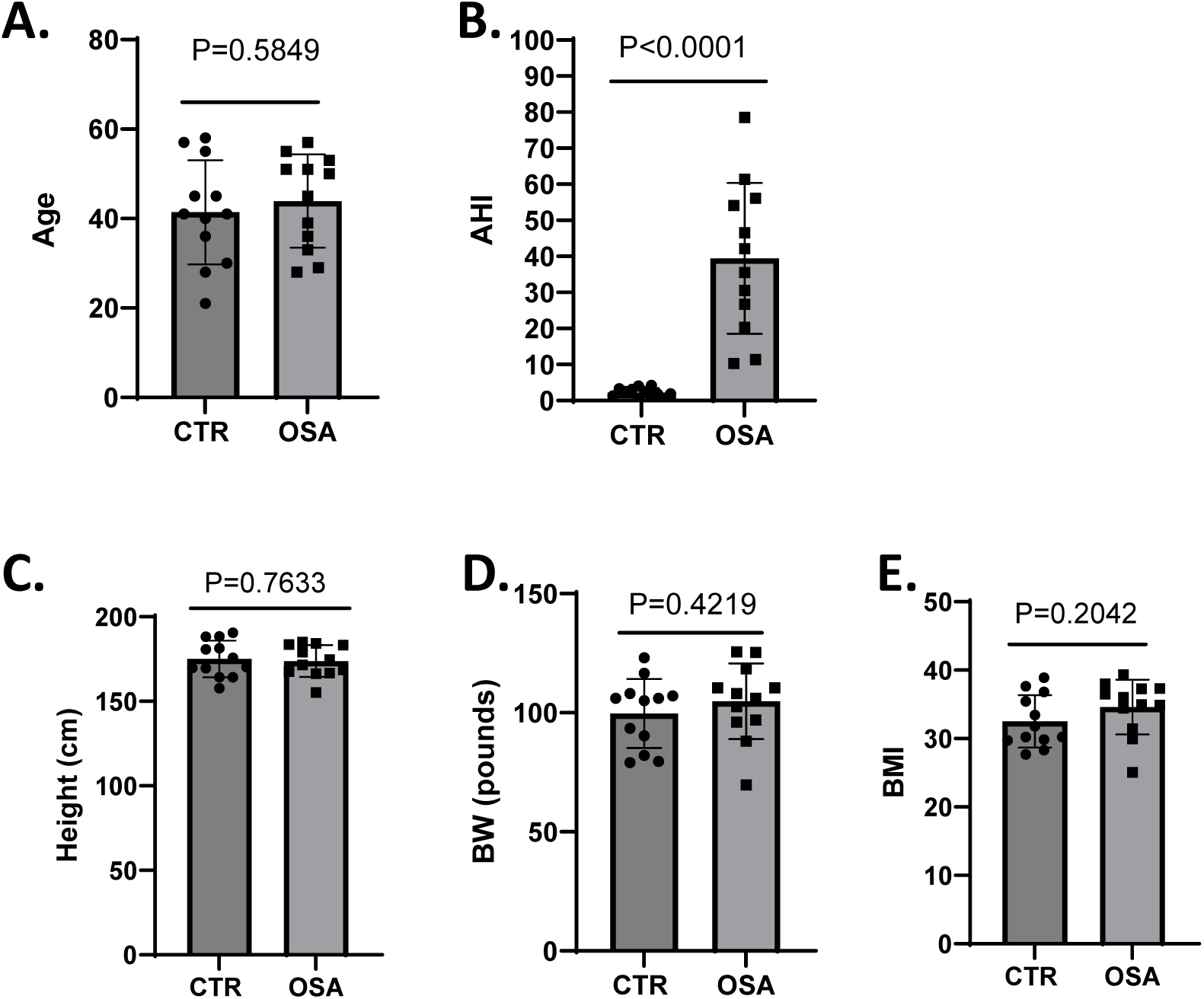
Demographic information of patients with obstructive sleep apnea and control subjects. A, age; B, AHI; C, Height; D, body weight; E, BMI. n=12 in each group. P values are indicated in each bar graph. Statistical analysis is conducted by student t test.

### Aldosterone content is not different in cell lysates between OSA and control subjects

Given that patients with obstructive sleep apnea (OSA) typically exhibit elevated blood pressure, our objective was to investigate whether adipocyte aldosterone levels are heightened in OSA patients and whether this could contribute to hypertension. To address this, we analyzed aldosterone levels in fat tissue lysates. Our findings showed no significant difference in aldosterone levels between subjects with OSA and the control group (see Fig. 2A). Subsequently, we examined the adipocyte protein expression of CYP11B2, a key enzyme involved in aldosterone synthesis, and similarly found no significant variation between OSA and control subjects (see Fig. 2B and C).

**Fig. 2.**
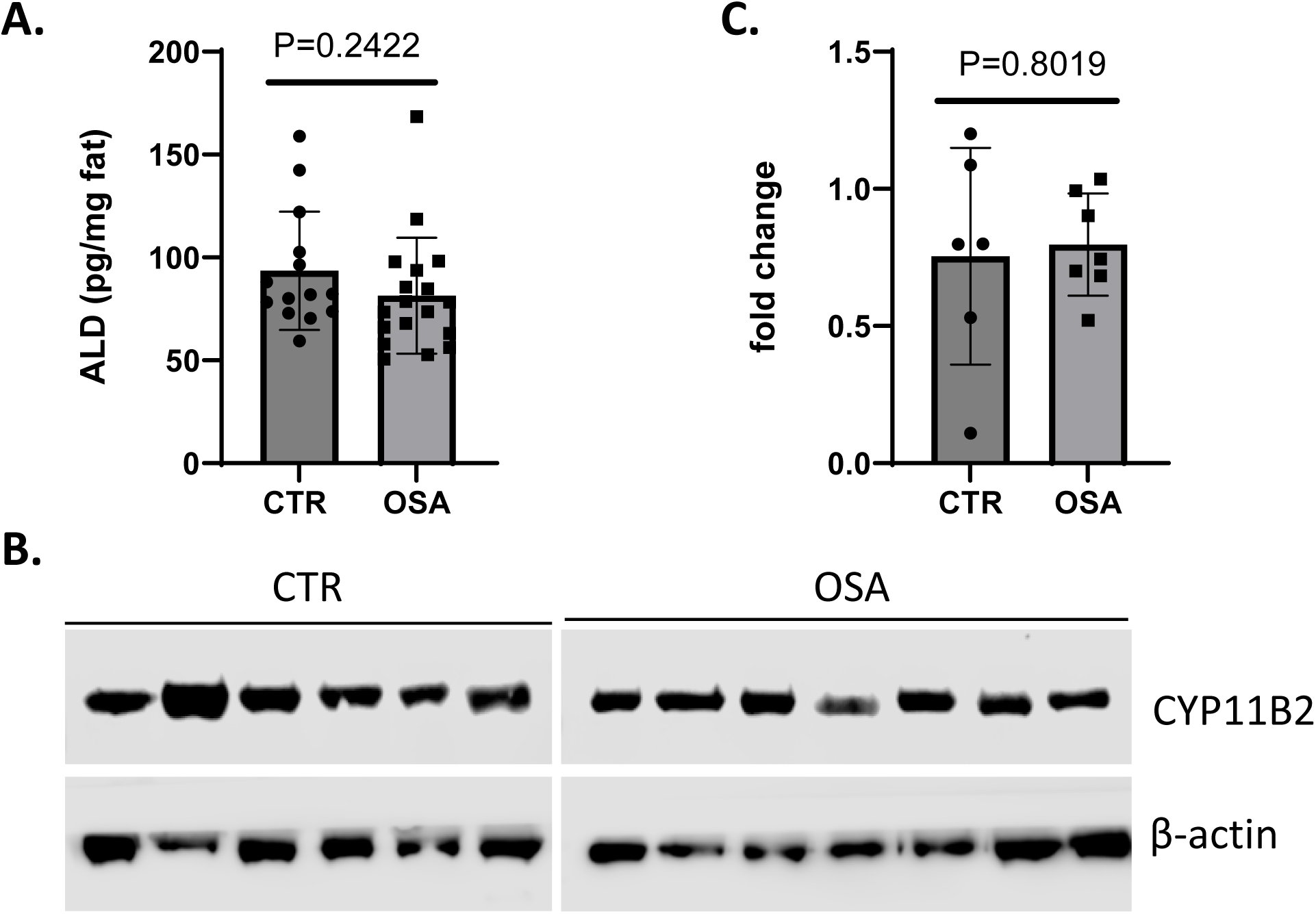
Aldosterone content and CYP11B2 expression in human fat tissues. Fat tissues from human were homogenized and protein concentrations were determined by BCA analysis. The cell lysates were subjected to ELISA assay for aldosterone content. Western blot was performed for the key gene expression of CYP11B2 responsible for ALD synthesis. A, ALD content in cell lysates; B. western blot analysis for CYP11B2 expression; C. quantification of CYP11B2. P values are indicated in the bar graphs.

### Aldosterone content is not different in cell lysates and conditioned medium between preadipocyte cultures exposed to normoxia and intermittent hypoxia

To complement our findings from patient samples, we employed cell culture methods to assess aldosterone production in human SVF preadipocytes derived from adipose tissues. As depicted in Fig 3A & 3B, aldosterone content exhibited no differences in the conditioned medium and cell lysates between normoxia and intermittent hypoxia (IH), Furthermore, western blot analysis revealed no significant differences in CYP11B2 protein expression under IH compared to normoxia conditions (Fig 3C & 3D), nor CYP11B2 mRNA levels as determined by qPCR analysis (data not shown).

**Fig. 3.**
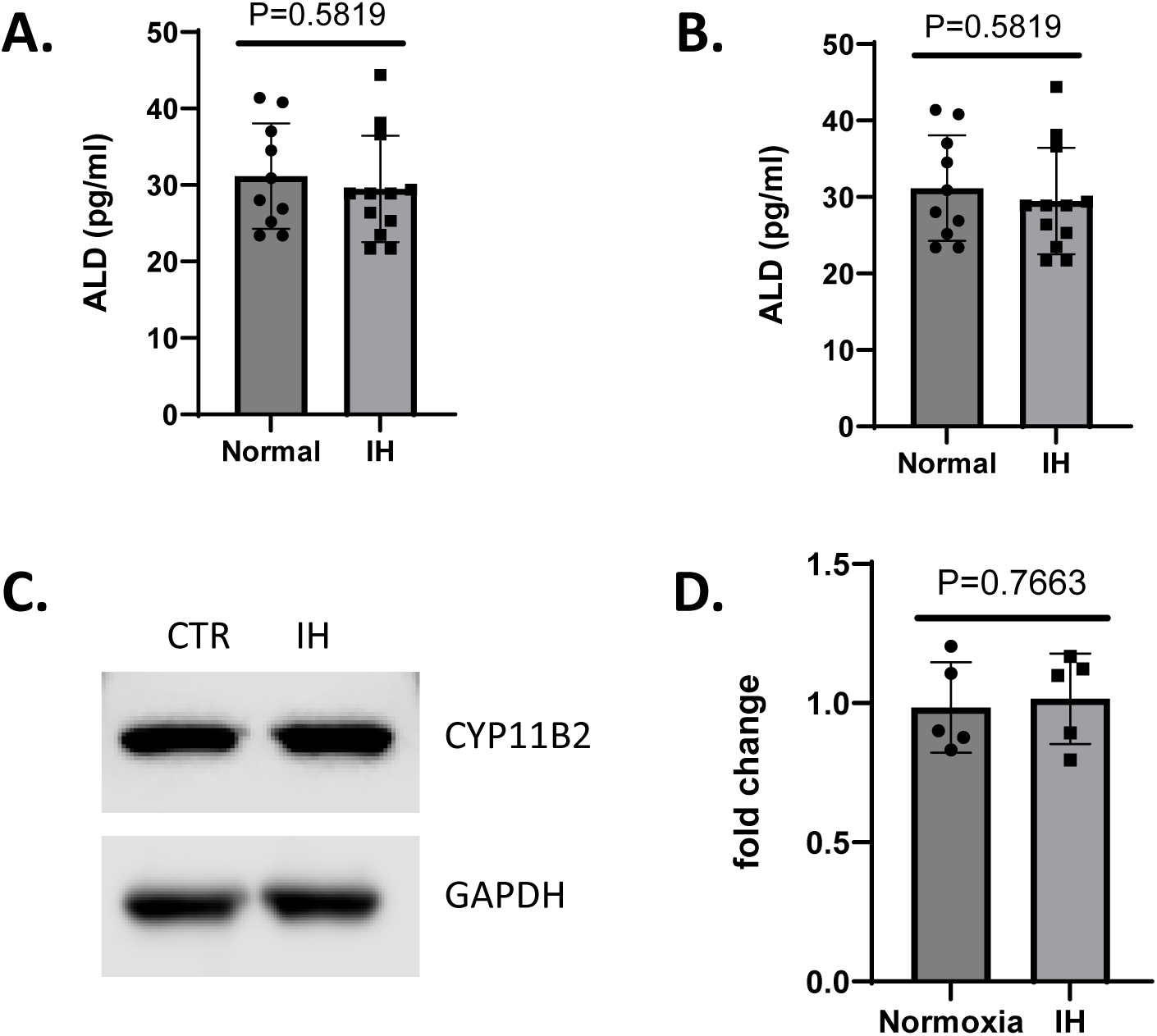
Aldosterone content and CYP11B2 expression in human preadipocytes. Subcutaneous abdominal fat tissues were isolated from human by liposuction. Strama vascular frictions (SVF) were isolated from the fat tissues and cultured in preadipocyte medium (Zenbio), followed by normoxia or intermittent hypoxia treatment for 7 days, and protein concentrations were determined by BCA analysis. The cell lysates and conditional medium were subjected to ELISA assay for aldosterone content. Western blot was performed for the key gene expression of CYP11B2 responsible for ALD synthesis. A, ALD content in conditional medium; B, ALD content in cell lysates; C. western blot analysis for CYP11B2 expression; D. Quantification of CYP11B2. P values are indicated in the bar graphs.

## Discussion

Obstructive sleep apnea (OSA) is a prevalent sleep disorder characterized by recurrent episodes of upper airway obstruction during sleep, leading to disrupted breathing patterns, intermittent hypoxia, sleep fragmentation, and daytime sleepiness. OSA has emerged as a significant public health concern due to its association with various cardiovascular and metabolic disorders, including hypertension and obesity (Jean-Louis *et al*., 2008). Understanding the complex interactions between OSA, hypertension, and obesity is essential for effective management and prevention of associated complications.

Hypertension, or high blood pressure, is a leading risk factor for cardiovascular disease and stroke, affecting millions of individuals worldwide. Studies have consistently demonstrated a strong association between OSA and hypertension, with estimates suggesting that up to 50-60% of patients with OSA have coexisting hypertension (Wolk *et al*., 2003; Narkiewicz *et al*., 2005; Golbin *et al*., 2008; Okcay *et al*., 2008). The pathophysiology underlying this relationship involves multiple mechanisms, including sympathetic activation, oxidative stress, inflammation, endothelial dysfunction, and altered autonomic control.

Furthermore, obesity is a well-established risk factor for both OSA and hypertension (Wolk *et al*., 2003; Xia *et al*., 2023). Excess adipose tissue, particularly in the upper body, can lead to mechanical narrowing of the upper airway, predisposing individuals to airway collapse during sleep. Additionally, adipose tissue secretes various bioactive molecules known as adipokines, which contribute to systemic inflammation, insulin resistance, dyslipidemia, and endothelial dysfunction, all of which can exacerbate cardiovascular risk factors.

Given the complex interactions between OSA, hypertension, and obesity, researchers have sought to identify potential mechanistic links and therapeutic targets to mitigate the adverse cardiovascular outcomes associated with these conditions. One such target of interest is aldosterone, a hormone primarily produced by the adrenal glands that plays a crucial role in regulating sodium and water balance, blood pressure, and electrolyte homeostasis (Awosika *et al*., 2023; Laffin *et al*., 2023; Pitt & Vaidya, 2023).

Aldosterone excess has been proposed as a key mechanism that could link obesity, OSA and hypertension, particularly in patients with resistant hypertension (Oparil *et al*., 2003; Calhoun & Harding, 2010; Clark *et al*., 2012; Siddiqui & Calhoun, 2017; Calhoun, 2018; Calhoun *et al*., 2019). Given the potential involvement of aldosterone in OSA-related hypertension and obesity, researchers have conducted studies to investigate its levels and activity in both systemic circulation and adipose tissue. However, the findings regarding the impact of OSA on aldosterone levels in adipose tissue have been limited and inconclusive (Briones *et al*., 2012; Wang *et al*., 2021).

To address these knowledge gaps, we conducted a comprehensive analysis of aldosterone specifically in adipose tissue from OSA patients. We hypothesized that OSA-induced intermittent hypoxia, may stimulate aldosterone secretion locally within adipose depots, thereby contributing to hypertension and metabolic dysregulation in these individuals.

Our study enrolled a diverse cohort of OSA patients, ranging from mild to severe cases, and carefully matched them with non-OSA controls based on age, sex, and BMI. We utilized state-of-the-art methodologies to obtain adipose tissue biopsies from subcutaneous fat depots to assess aldosterone levels and activity.

In contradiction to our original hypothesis, our data revealed no significant differences in adipose tissue aldosterone levels between OSA patients and controls. Furthermore, even exposure of preadipocytes to intermittent hypoxia *in vitro* did not result in any increase in the crucial enzyme CYP11B2 expression nor in aldosterone production.

Despite the lack of direct association between OSA and aldosterone levels in adipose tissue in our study, it is essential to recognize that hypertension and obesity in OSA patients are multifactorial conditions influenced by a complex interplay of genetic, environmental, and physiological factors (Abbasi *et al*., 2021). Our data confirm that adipose tissue is indeed a source of aldosterone production, so that increasing levels of obesity would be expected to result in increased levels of aldosterone. Indeed, plasma aldosterone level is associated with BMI in cohorts of patients with hypertension (Mule *et al*., 2008; Rossi *et al*., 2008). However, our study suggests that the comorbidity of OSA does not further potentiate adipose tissue aldosterone production. However, our findings apply only to subcutaneous adipose tissue and do not exclude the possibility that visceral fat aldosterone production may be impacted in patients with OSA.

In conclusion, our data suggest that aldosterone production in adipose tissue is not be potentiated by IH nor by OSA.

## Additional Information

### Competing interesting

VKS is a consultant for Axsome, Jazz Pharmaceuticals, Lilly, and ApniMed and serves on the Sleep Number Scientific Advisory Board. No conflicts of interests to disclosure.

### Author contributions

YD and VKS conceptualized the experiments. YD carried out the aldosterone content analysis, measured CYP11B2 expression and conducted the data analysis. YD wrote the manuscript, which was subsequently refined by VKS and other coauthors. JB, SK and PS facilitated the collection of demographic data and conducted human fat tissue biopsies. PS provided valuable comments during manuscript preparation.

### Funding

This research was supported by NIH grant HL 65176 to VKS and PS.

## Acknowledgements

We express our gratitude to the participants who generously contribute subcutaneous abdominal fat tissue samples for this study. Additionally, we extend our sincere appreciation to Dr. Zvonimir Katusic and Dr. Tongrong He for their technical support. Special thanks are also due to Debra Pfeifer and Jan Bukartyk for their assistance.

## Supporting information

Additional supporting information can be found online regarding participants and SVF isolation and preadipocyte culture from human subcutaneous abdominal fat tissues.

## Notes

### Competing Interest Statement

The authors have declared no competing interest.

### Summary of Updates

Due to an error, a human MC# was mistakenly included in the supplementary materials. In accordance with Mayo Clinic policy, human medical record numbers (MC#s) must not be disclosed or included in public documents, even inadvertently. To comply with this policy and safeguard confidentiality, we will remove the MC# from the supplementary materials. This ensures adherence to institutional standards for privacy and data security. We apologize for this oversight and are taking the necessary steps to rectify it promptly. Please let us know if further clarification or actions are required regarding this revision.

