## Supplementary material for "Adipocyte Aldosterone in Obstructive Sleep Apnea": supple figure

**Supple Figure 1**. Isolated SVF and cultured in preadipocyte medium

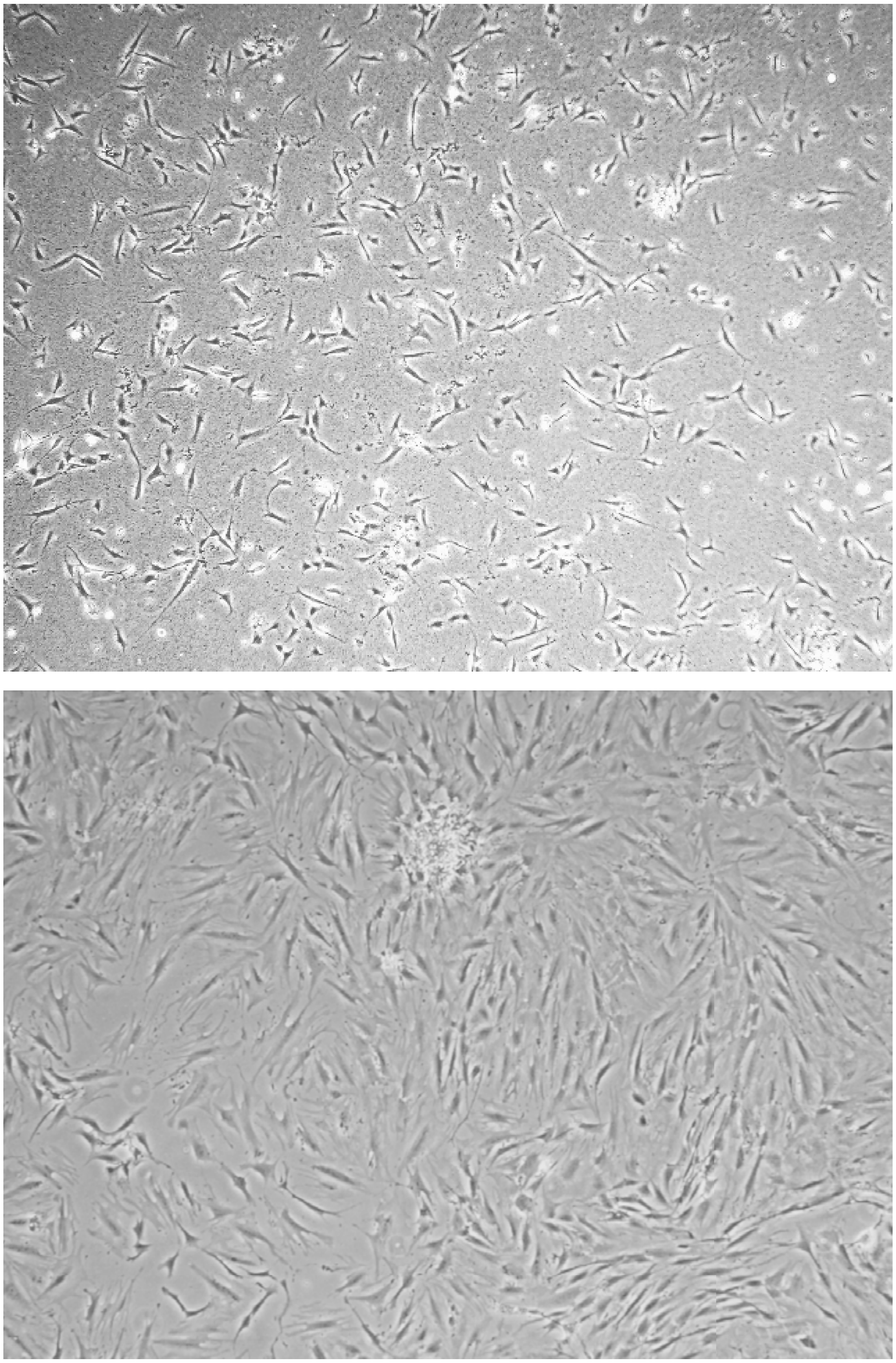

4 day cultured preadipocytes P0 (4x)

8 day cultured preadipocytes P1 (4x)
